# A little longer, a lot better: simulation-guided exploration of extended-length single-end barcoded reads for structural variant detection

**DOI:** 10.1101/2025.03.31.646392

**Authors:** Can Luo, Yichen Henry Liu, Han Liu, Zhenmiao Zhang, Lu Zhang, Brock A. Peters, Xin Maizie Zhou

## Abstract

Accurate detection of genetic variants, including single nucleotide polymorphisms (SNPs), small insertions and deletions (INDELs), and structural variants (SVs), is essential for comprehensive genomic analysis. While short-read sequencing performs well for SNP and INDEL detection, it remains limited in resolving SVs, particularly in complex genomic regions, due to its short read length. Linked-read sequencing technologies, such as single-tube Long Fragment Read (stLFR), partially address this limitation by incorporating molecular barcodes to provide long-range information. In this study, we evaluate conventional paired-end linked reads (PE100_stLFR) and explore a conceptual extension: long single-end barcoded reads of 500 bp (SE500_stLFR) and 1000 bp (SE1000_stLFR). We developed_stLFR-sim, a Python-based simulator that reproduces the_stLFR workflow and enables realistic benchmarking. Using a high-quality T2T assembly of HG002, we generated multiple datasets across 12 sequencing configurations. SVs were called using Aquila_stLFR (v2) and benchmarked against the Genome in a Bottle (GIAB) HG002 SV truth set with Truvari. We show that simulated PE100_stLFR closely matches real data, validating the simulation framework. Increasing read length consistently improves SV detection accuracy, with SE1000_stLFR achieving the best performance and approaching long-read methods while outperforming short-read and pangenome-based approaches. Collectively, our results highlight the strong potential of long single-end barcoded reads for improving SV detection, and suggest that even modest increases in read length, when combined with barcode information, can provide a cost-effective and practical strategy for enhancing future sequencing technologies and SV discovery.

## Background

Advancements in sequencing technologies have revolutionized genomics study by enabling precise and high-throughput analysis of DNA, offering unprecedented opportunities to understand genetic variation [1], disease mechanisms [2], and evolutionary biology [3]. Among these advancements, short-read sequencing [4, 5] has become the backbone of many genomic studies due to its cost-effectiveness, scalability, and high accuracy in detecting small variants, such as single nucleotide polymorphisms (SNPs) and small insertions and deletions (indels). However, despite its transformative impact, the intrinsic limitations of short-read sequencing in terms of read length impose significant challenges when addressing more complex genomic features, such as structural variants (SVs), repetitive regions, and chromosomal rearrangements. The inability to span long repetitive sequences or phase variants across distant loci restricts its utility for comprehensive genome characterization, leaving gaps in our understanding of genetic variation.

To overcome these challenges, innovative technologies have been developed to extend the capabilities of sequencing, providing long-range information and improved resolution of complex genomic features. One notable advancement is linked-read sequencing, which combines short-read sequencing with molecular barcoding to retain long-range context [6,7]. This method starts by fragmenting high molecular weight (HMW) DNA into long segments, typically tens to hundreds of kilobases in length. Each fragment is encapsulated into microfluidic droplets or physically separated into individual compartments, where it undergoes amplification and tagging with a unique molecular barcode. Importantly, all short reads derived from a single DNA fragment carry the same unique barcode, allowing them to be computationally grouped and assigned to their original long DNA molecule. Once the barcoded DNA fragments are sequenced, bioinformatics tools can reconstruct the long-range structure of the original DNA by clustering reads based on their shared barcode [8–10]. This enables linked-read sequencing to effectively span long regions of the genome, even with the relatively short read lengths produced by sequencing platforms like Illumina. This methodology enhances genome assembly, variant detection, and haplotype phasing by preserving the positional information of DNA fragments, which is critical for resolving structural variants and complex regions [11–15].

Among linked-read technologies, the 10x Genomics Chromium platform stands out as a pioneering implementation [6]. It uses microfluidics to partition HMW DNA into thousands of droplets, each containing a unique barcode and reagents for DNA amplification. This approach ensures high efficiency and scalability, enabling the analysis of large and complex genomes. However, one limitation of this method is the occurrence of multiple DNA molecules within a single droplet, leading to potential barcode collisions and reduced resolution in some cases. The advent of single-tube long fragment read (stLFR) technology addressed this issue by eliminating the need for droplet-based separation and ensuring a “near” one-to-one correspondence between individual DNA molecules and barcodes [7]. Using a bead-based chemistry, stLFR introduces barcodes directly to DNA fragments in solution, simplifying the workflow and improving the accuracy of molecule-to-barcode association. This innovation further enhances the resolution of linked-read sequencing, particularly for applications requiring high precision in structural variant detection and haplotype phasing. Despite these advancements, the SV discovery performance of traditional paired-end linked reads still remains inferior to that of long-read sequencing. This led us to ask whether modest modifications, such as slightly extending the read length, combined with the power of barcode information could achieve performance comparable to more expensive long-read technologies.

Motivated by this idea, this study explores a conceptual extension of linked-read technology: long singleend barcoded reads with lengths of 500 bp and 1000 bp. Unlike the traditional paired-end reads commonly used in linked-read sequencing, barcoded single-end reads could simplify library preparation workflows while leveraging longer individual reads per barcode to provide improved mapping resolution. This design has the potential to enhance structural variant (SV) detection, particularly in genomic regions that are difficult to resolve using conventional short-read and paired-end approaches. To evaluate this concept, we simulated different linked-read data, including conventional paired-end linked read(PE100_stLFR) and the conceptual single-end linked-read(SE500_stLFR and SE1000_stLFR) of 12 sequencing configurations(Table 1), respectively. We first compared the variant calling performance between simulated PE100_stLFR data and real PE100_stLFR data to verify that our simulation framework generates realistic linked-read datasets. We then systematically assessed SV detection performance using stLFR-sim–generated libraries, including both conventional paired-end linked reads and conceptual single-end barcoded reads. Our results demonstrate that barcoded sequencing with moderately extended read lengths can substantially improve SV detection accuracy, achieving performance approaching that of long-read sequencing and pangenome-based short-read genotyping strategies.

**Table 1.** The table summarizes the simulation settings for all experiments (datasets). All datasets share the same *N*_*FL*_ (number of fragments per barcode), insert size, and error rate. PE100_stLFR, SE500_stLFR, and SE1000_stLFR indicate the read length and whether sequencing is paired-end or single-end. Experiments marked as 35x correspond to sequencing depths of 35x. Experiments (EXP) 1-12 are divided into three groups: 1-4, 5-8, and 9-12. Across these groups, *µ*_*FL*_ varies, while *C*_*F*_ and *C*_*R*_ vary within each group.

| Shared Parameters: $N_{FL}=1$; Insert Size=600 bp; Error Rate=1% | | | |
| --- | --- | --- | --- |
|  | PE100_stLFR/SE500_stLFR/SE1000_stLFR 35x |  |  |
| | $C_F$ | $C_R$ | $\mu_{FL}(kb)$ |
| EXP1 | 350 | 0.1 | 50 |
| EXP2 | 140 | 0.25 | 50 |
| EXP3 | 70 | 0.5 | 50 |
| EXP4 | 47 | 0.75 | 50 |
| EXP5 | 350 | 0.1 | 75 |
| EXP6 | 140 | 0.25 | 75 |
| EXP7 | 70 | 0.5 | 75 |
| EXP8 | 47 | 0.75 | 75 |
| EXP9 | 350 | 0.1 | 100 |
| EXP10 | 140 | 0.25 | 100 |
| EXP11 | 70 | 0.5 | 100 |
| EXP12 | 47 | 0.75 | 100 |

These findings suggest that if long single-end barcoded reads become technically feasible, it could provide a highly cost-effective strategy for accurate structural variant discovery. Our work therefore offers a promising and practical blueprint for the future design of linked-read sequencing technologies.

## Methods

### Simulator pipeline

In this study, we present stLFR-sim, a Python-based simulator designed to generate linked-read sequencing data that mimics the output of the stLFR platform and Illumina sequencers. While it draws partial inspiration from the methodology of LRTK-sim [16], stLFR-sim is an original tool that introduces several novel features. Compared to LRTK-sim which is a linked-read simulator for 10X Chromium System, stLFR-sim is optmized for stLFR linked-reads simulation. Furthermore, stLFR-sim offers the original feature of simulating long single-end barcoded reads. In detail, the simulator reproduces the stLFR linked-read sequencing workflow through four main steps: (1) generating a diploid reference genome, (2) simulating long DNA fragments, (3) assigning barcodes to DNA fragments, and (4) generating barcoded Illumina short reads. This pipeline enables realistic simulation of linked-read data, making stLFR-sim a valuable resource for benchmarking sequencing workflows and downstream analysis tools. Each step is elaborated below to provide a comprehensive overview of the simulation process.

#### 1. Generating the Diploid Reference Genome

stLFR-sim begins with a sample-specific diploid genome as input. While the original LRTK-sim pipeline generated this by inserting phased, highconfidence variants into a reference genome, our approach instead utilizes a phased, realistic diploid assembly from the HG002 sample (see section titled “T2T Realistic Simulation” for details). This diploid reference serves as the foundation for generating synthetic linked-read data, allowing the simulation to capture the genomic complexity of a real individual.

#### 2. Simulating Long DNA Fragments

With the diploid genome defined, stLFR-sim proceeds to simulate long DNA fragments. The physical coverage, denoted *C*_*F*_, represents the cumulative coverage of all DNA fragments across the genome. The total amount of DNA, *V*, is calculated as *V* = *C*_*F*_ *× L*, where *L* is the length of the reference genome. To model both haplotypes equally, each is assigned half of the total physical coverage, i.e., *C*_*F*_ */*2. Fragment start positions are randomly distributed across the genome, and fragment lengths are drawn from an exponential distribution with a mean of *µ*_*FL*_. This step ensures that the simulated fragment size distribution closely mirrors that observed in real linked-read sequencing data.

#### 3. Barcoding DNA Fragments

In this step, stLFR-sim emulates the barcoding process used in stLFR technology. There are originally 1536 unique 10-mer barcodes in the stLFR barcode list. stLFR-sim randomly selects 3 barcodes from the list, thereby generating up to 3.62 billion unique 30-mer barcode sequences for labeling DNA fragments. This number (3.62 billion) represents the theoretical maximum number of barcodes; however, the actual number utilized depends on the values of *C*_*F*_ and *µ*_*FL*_, and is typically much lower than the maximum. Specifically, the actual number of barcodes can be estimated as *N*_*bc*_ = *C*_*F*_ *×* genome size / *µ*_*FL*_. stLFR-sim generates “pseudo” partitions, each assigned a unique barcode. In contrast to LRTK-sim’s pipeline, which simulates the stochastic barcoding process by assigning multiple fragments to a single partition, stLFR-sim reflects the design of stLFR technology, which enables precise assignment of one fragment per partition. Consequently, each DNA fragment in stLFR-sim is assigned to a distinct partition with a unique barcode, accurately modeling the one-fragment-per-barcode characteristic of stLFR.

#### 4. Simulating Barcoded Illumina Short Reads

The final step in stLFR-sim involves generating barcoded Illumina short reads to cover the simulated long DNA fragments. The total number of reads is calculated as 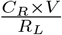, where *C*_*R*_ denotes the desired read coverage, *V* is the total DNA volume, and *R*_*L*_ is the read length, typically set to 150 bp by default. Reads are uniformly distributed across each fragment to mimic even sequencing coverage.

Paired-end short reads are generated with insert sizes sampled from a normal distribution centered at a userdefined mean *µ*_*IS*_. Short reads are tagged with barcodes derived from their associated DNA fragments. To further enhance realism, stLFR-sim incorporates empirical base quality profiles derived from linked-read datasets sequenced on an Illumina HiSeq X platform. Sequencing errors are introduced at random positions based on these quality scores, providing a more accurate simulation of real-world sequencing data.

#### 5. Simulating Other Sequencing Data Types

To broaden the scope of the simulation, stLFR-sim also supports the generation of standard Illumina short reads (without barcodes) and long single-end barcoded reads. For simulating unbarcoded Illumina short reads, the barcoding step is simply disabled while the rest of the pipeline remains unchanged. To simulate long single-end barcoded reads, the paired-end sequencing module is replaced with a single-end mode, which randomly samples fixed-length reads from each fragment.

In summary, stLFR-sim provides several advantages over existing linked-read simulators. First, it allows users to flexibly configure key parameters, including *C*_*F*_, *C*_*R*_, *µ*_*FL*_, *R*_*L*_, and *µ*_*IS*_, to accurately model specific experimental conditions. Second, we have extended its functionality to support the simulation of standard Illumina short reads (without barcodes) and long single-end barcoded reads, enhancing its versatility. Finally, stLFR-sim is a lightweight, all-in-one Python package that requires no third-party software. In contrast to simulators like LRSIM, which depend on multiple external tools, stLFR-sim is self-contained and capable of simulating multiple libraries in a single run, making it both efficient and user-friendly for exploring a wide range of experimental designs.

### T2T realistic simulation

To simulate reads sequenced within realistic genomic contexts, we utilized the phased diploid assembly of HG002 released by the Human Pangenome Reference Consortium (HPRC) [17]. The paternal and maternal haplotype assemblies were independently aligned to the T2T-CHM13(v2).0 reference using minimap2. Contigs from each haplotype that aligned to the target chromosome (e.g., chromosome 6) were then extracted using SAMtools and used as input for our simulation.

~~~
minimap 2 -- secondary =no -a -- eqx -Y -
   x asm20 -s 200000 -z 10000,50 -r
   50000 -- end - bonus =100 -O 5,56 -E
   4,1 -B 5 ${ ref} ${ asm } | samtools
   sort -o ${ bam }
~~~

### Simulation configurations

Read libraries were simulated under various configurations to systematically investigate the impact of different parameter combinations. PE100 stLFR libraries consist of paired-end reads with a length of 100 bp, while SE500 stLFR and SE1000_stLFR libraries contain single-end barcoded reads with lengths of 500 bp and 1000 bp, respectively. A total of 12 simulation experiment (EXP1-EXP12) were conducted for each library type. These experiments were categorized into three groups: EXP1-EXP4, EXP5-EXP8, and EXP9-EXP12. Within each group, the parameters *C*_*F*_ and *C*_*R*_ were varied, while the parameter *µ*_*FL*_ differed between groups. All groups shared consistent values for insertion size, error rate, and number of fragments per barcode (*N*_*FL*_).

More specifically, the *C*_*F*_ - *C*_*R*_ combinations ranged from 350-0.1 to 47-0.75 within each group. The *µ*_*FL*_ values were set to 50 kb, 75kb, and 100 kb for EXP1-EXP4, EXP5-EXP8, and EXP9-EXP12, respectively. Across all experiments, the insertion size, error rate, and number of fragments per barcode (*N*_*FL*_) were consistently set to 600 bp, 1%, and 1, respectively. Full simulation configurations are summarized in Table 1.

### Aquila_stLFR (v2) and SV calling

We developed Aquila stLFR (v2) as an updated pipeline for structural variant (SV) calling, based on the original Aquila stLFR framework [9]. Compared to the previous version, it offers the new feature of processing long single-end barcoded reads while having more optimized software structure. This version is a reference-assisted tool designed to perform local *de novo* assembly of stLFR linked-read data. It reconstructs haplotype-specific blocks using barcode-tagged long fragment reads (LFRs) and performs localized assembly within small genomic regions to achieve high contiguity and accurate SV detection.

Aquila stLFR (v2) processes stLFR data in three main steps: (1) Haplotype phasing: Long fragment reads (LFRs) are reconstructed and partitioned into haplotype-specific blocks using barcode information and heterozygous SNPs. (2) Local assembly: Within 100 kb genomic chunks, reads are locally assembled using SPAdes. Resulting mini-contigs are then iteratively merged to form full-length haplotype-resolved contigs. (3) SV detection: Structural variants are identified by aligning haploid contigs to the reference genome using Minimap2 [18], followed by a custom SV detection pipeline based on VolcanoSV-vc [19].

While steps 1 and 2 remain unchanged from the original Aquila stLFR, step 3 has been enhanced in version 2 to incorporate the VolcanoSV-vc methodology for more robust SV detection.

VolcanoSV-vc is the variant calling module of VolcanoSV [19], originally designed for long-read sequencing data. For our linked-reads application, we adapted VolcanoSV-vc by disabling the long-read-specific signature scanning module. In our workflow, VolcanoSV-vc detects SVs from haplotype-resolved contigs aligned to the reference genome using Minimap2. SVs, such as large insertion and deletion SVs, are identified based on CIGAR string information and split alignment signatures.

SV signatures are collected independently for each haplotype, clustered using a nearest-neighbor algorithm, and refined to eliminate redundancy. Within each cluster, the longest signature is retained as the final SV call. Genotypes are then assigned based on the presence of SVs across haplotypes: “1|1” if present in both, and “0|1” or “1|0” if present in only one. Despite potential assembly artifacts, VolcanoSV-vc uses empirically optimized thresholds to ensure accurate clustering and genotyping in the linked-reads context.

The following commands were used to generate the SV calls.

~~~
Aquila_stLFR 2 _step 1 . py \
    --fastq_file ${ reads_fastq } \
    --bam_file ${ bamfile } \
    --vcf_file ${ SNP_vcf} \
    --sample_name HG002 \
    --out_dir ${ outdir} \
    --uniq_map_dir ./ Aquila_stLFR /
       Uniqness_map_hg 19 \
    --chr_start ${ chr_num } -- chr_end
        ${ chr_num } -- num_threads $t -
       r $ dtype
Aquila_stLFR 2 _step 2 . py \
    --out_dir ${ outdir} \
    --num_threads $t \
    --reference $ reference \
    --chr_start ${ chr_num } -- chr_end
       ${ chr_num } -r $ dtype
Aquila_stLFR 2 _step 3 . py \
         -contig ${ outdir }/
             Assembly_Contigs_files /
             Aquila_Contig_chr ${ chr_num
             }. fasta \
         -ref ${ reference } \
         -o ${ outdir }/ SV_ouput/ \
         -chr ${ chr_num } -t $t
~~~

### SNP and small INDEL calling

For SNP and INDEL calling, simulated reads were first aligned to the reference genome using BWA-MEM (v0.7.17) [20] for long single-end barcoded reads, and EMA (v0.6.2) [21] for stLFR paired-end reads. Read group information was then added to the alignments using Picard. Finally, SNP and INDEL calling was performed using the GATK (v4.3.0) pipeline [22].

The following commands were used to generate the SNP and INDEL calls:

~~~
bwa mem \
    ${ reference } \
    ${ R 1 _fastq } \
    ${ R 2 _fastq } \
    -t 30 |samtools sort -o ${ bwa_bam
       }
ema align \
    -1 ${ R 1 _fastq } \
    -2 ${ R 2 _fastq } \
    -r ${ reference } -t 30 |samtools 
       sort -@ 30 -o ${ ema_bam }
java - jar picard . jar 
    Add OrReplace Read Groups \
     I=${ raw_bamfile } \
     O=${ bamfile } \
     RGID =4 \
     RGLB = lib1 \
     RGPL = ILLUMINA \
     RGPU = unit1 \
     RGSM =20
gatk Haplotype Caller -I $ bamfile -O 
    gatk . vcf -R ${ reference }
~~~

### Benchmarking

For SV detection evaluation, we used Truvari (v4.0.0) [23] to benchmark our SV callsets against the Genome in a Bottle (GIAB) HG002 SV Tier1 v0.6 highconfidence truth set [24] on the hg19 reference genome. Truvari was run with a maximum reference distance of 500 bp (*refdist*), a minimum allele sequence similarity of 50% (*pctsim*), a minimum allele size similarity of 50% (*pctsize*), a minimum reciprocal overlap of 1% (*pctovl*), and restrictions to high-confidence regions (*includebed*) and PASS variants (*passonly*). All other parameters were left at their default values.

To contextualize the performance of our method, we additionally compared it with representative approaches from three major paradigms of SV analysis: conventional short-read SV calling, pangenome-based short-read genotyping, and long-read–based SV detection. Specifically, Manta (v1.6.0) [25] was used as a representative short-read SV caller, PanGenie (v4.2.1) [26] as a pangenome-based short-read genotyper, and VolcanoSV [19] as a long-read–based SV caller. Manta and PanGenie were both applied to the same Illumina short-read dataset. For PanGenie, genotyping was performed using a pangenome graph constructed from nine Human Pangenome Reference Consortium (HPRC) samples: HG01109, HG01243, HG02055, HG02080, HG02109, HG02145, HG02723, HG03098, and HG03492. VolcanoSV was applied to PacBio HiFi reads. Unless otherwise noted, each method was run using its recommended workflow and standard settings, and the resulting callsets were benchmarked within the same Truvari-based evaluation framework.

SNP and small INDEL benchmarking was performed using hap.py against the NIST v4.2.1 HG002 GRCh37 call set on the hg19 reference genome. Performance was assessed in both high-confidence and difficult genomic regions, including the major histocompatibility complex (MHC), tandem and homopolymer repeats (tandemRep), segmental duplications (segdup), and low-mappability regions (lowmap), using the corresponding BED files from the NIST v4.2.1 genome stratifications. Default parameters were used for all hap.py evaluations.

## Results

Our goal was to explore the potential of linked reads for structural variant (SV) detection and to investigate how sequencing configurations influence SV calling performance. We were also interested in evaluating the potential of long single-end barcoded reads. Therefore, using the simulator stLFR-sim and a high-quality diploid assembly of sample HG002 as the reference genome, we simulated a series of stLFR linked-read datasets, including PE100_stLFR, SE500_stLFR, and SE1000_stLFR. In stLFR-sim, multiple sequencing parameters are incorporated to generate realistic simulations, including *C*_*F*_ (coverage of long fragments), *C*_*R*_ (coverage of short reads on each long fragment), *µ*_*FL*_ (average fragment length), and sequencing error rate. By tuning these parameters, we were able to model a range of sequencing scenarios observed in real-world experiments. For each sequencing type, we designed 12 parameter configurations, which are detailed in Table 1. For each simulated paired-end library, reads were aligned using the barcode-aware aligner EMA (v0.6.2) [21]. For each simulated single-end library, reads were aligned using the aligner BWA-MEM (v0.7.17) [20]. SNPs were called using GATK (v4.3.0) [22], and SVs were detected using Aquila stLFR (v2) [9]. The resulting SV calls were then compared to the GIAB SV benchmark set [27] using the benchmarking tool Truvari (v4.0.0) [23], generating a comprehensive set of evaluation metrics. We further compared the SV calling performance of long single-end barcoded reads (analyzed using Aquila stLFR (v2)) with that of traditional paired-end short reads (analyzed using Manta (v1.6.0) [25] and PanGenie (v4.2.1) [26]) and long reads (analyzed using VolcanoSV [19]), demonstrating the strong potential of long single-end barcoded reads for accurate SV detection. Detailed analyses of these results are presented in the following sections.

### stLFR-sim reproduces real-data performance with high fidelity

In this session, we compared the sequencing configuration and variant calling performance between our simulated data and real data. In our simulation experiment, we simulated PE100_stLFR data across 12 different simulation configurations (Table 1). In our simulation pipeline, the determining parameters were *C*_*F*_ (coverage of long fragment), *C*_*R*_ (coverage of short reads on each long fragment), *µ*_*FL*_ (average fragment length), and sequencing error rate. Specifically, we simulated 12 different libraries for PE100_stLFR, among which *C*_*F*_ ranged from 47-350, *C*_*R*_ ranged from 0.1-0.75, *µ*_*FL*_ ranged from 50-100 kb. The error rate was set to 1% for all 12 libraries. For the real PE100_stLFR data, *C*_*F*_ was 931, *C*_*R*_ was 0.11, *µ*_*FL*_ was 13.9kb, and the error rate was 1%.

For SV calling, we first compared the overall simulated data across experiments and real PE100_stLFR data (Figure 1). We also compared the performance between simulated experiment 1 of PE100_stLFR and real PE100_stLFR (Table 2) since their sequencing configurations were most similar. From both analyses, the simulated and real PE100_stLFR showed an overall similar trend and accuracy: for insertion SV, the precision was high but the recall was relatively lower, while for deletion SVs, the recall was high but the precision was generally lower. A key objective of our simulation was to assess the potential performance gains of stLFR reads under configurations optimized for structural variant (SV) detection. To this end, we simulated fragment lengths ranging from 50 to 100 kb, exceeding the typical fragment lengths observed in real datasets (e.g., the real dataset used in this study had an average fragment length of 13.9 kb). As a result, the simulated PE100_stLFR dataset achieved a higher average F1 score than the real PE100_stLFR data, consistent with the expectation that longer fragments enhance SV-calling performance.

**Table 2.**
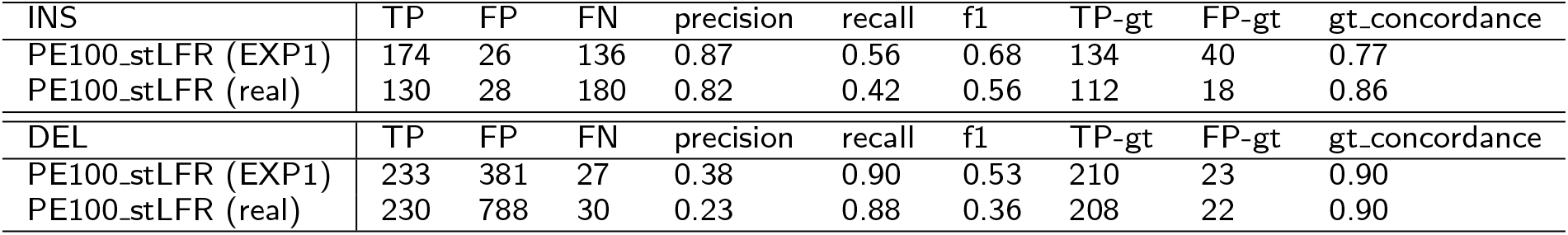
Comparison of structural variant calling performance for deletion and insertion SVs between PE100_stLFR (EXP1) and PE100_stLFR (real). Performance is summarized using true positives (TP), false positives (FP), false negatives (FN), precision, recall, F1 score, genotype true positives (TP-gt), genotype false positives (FP-gt), and genotype concordance.

| INS | TP | FP | FN | precision | recall | f1 | TP-gt | FP-gt | gt_concordance |
| --- | --- | --- | --- | --- | --- | --- | --- | --- | --- |
| PE100_stLFR (EXP1) | 174 | 26 | 136 | 0.87 | 0.56 | 0.68 | 134 | 40 | 0.77 |
| PE100_stLFR (real) | 130 | 28 | 180 | 0.82 | 0.42 | 0.56 | 112 | 18 | 0.86 |
| DEL | TP | FP | FN | precision | recall | f1 | TP-gt | FP-gt | gt_concordance |
| PE100_stLFR (EXP1) | 233 | 381 | 27 | 0.38 | 0.90 | 0.53 | 210 | 23 | 0.90 |
| PE100_stLFR (real) | 230 | 788 | 30 | 0.23 | 0.88 | 0.36 | 208 | 22 | 0.90 |

**Figure 1.**
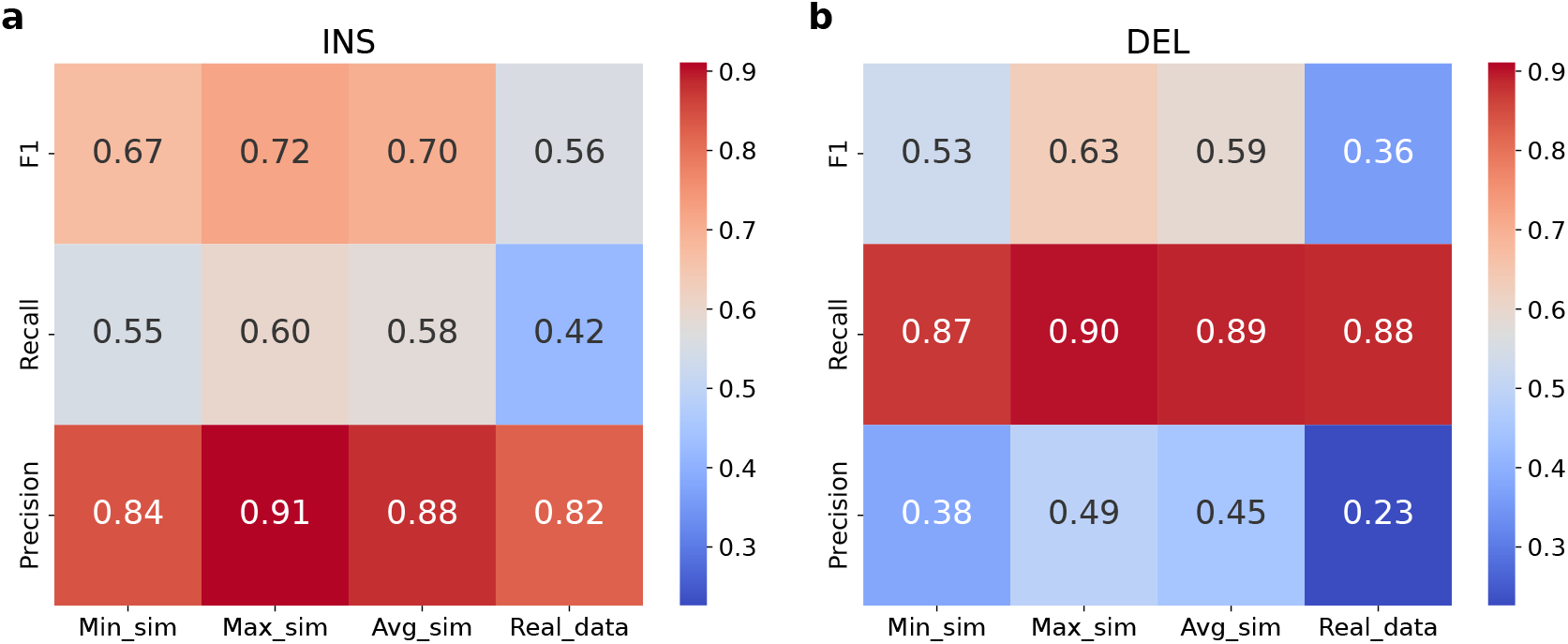
Simulated and real PE100_stLFR SV calling performance comparison. *T ruvari* (v4.0.0) was applied to benchmark the SV callset against the GIAB SV benchmark set. Across 12 different simulation configurations, the minimum (Min sim), maximum(Max sim) and average(Avg sim) value of F1, recall and precision score were calculated. **a**. F1, recall, precision for insertion SV calling. **b**. F1, recall, precision for deletion SV calling.

We also compared SNP, small INDEL calling performance between simulated and real PE100_stLFR data. For SNP and small INDEL calling (Figure 2), we evaluated performance across 12 different simulation configurations. Across each configuration, the minimum (Min sim), maximum (Max sim), and average (Avg sim) values of F1 score, recall, and precision were calculated. The evaluation was performed across multiple genomic regions, including all difficult regions (alldiff), low mappability regions (lowmap), segmental duplication regions (segdups), tandem repeat regions (tandemRep), the major histocompatibility complex (MHC), and the GIAB high-confidence regions for HG002 (highconf). Our simulated data on average showed comparable result to real data. Especially, in the high-confidence region, the difference between average simulation data F1 and real data F1 was only 0.01 for both SNP and small INDEL calling.

**Figure 2.**
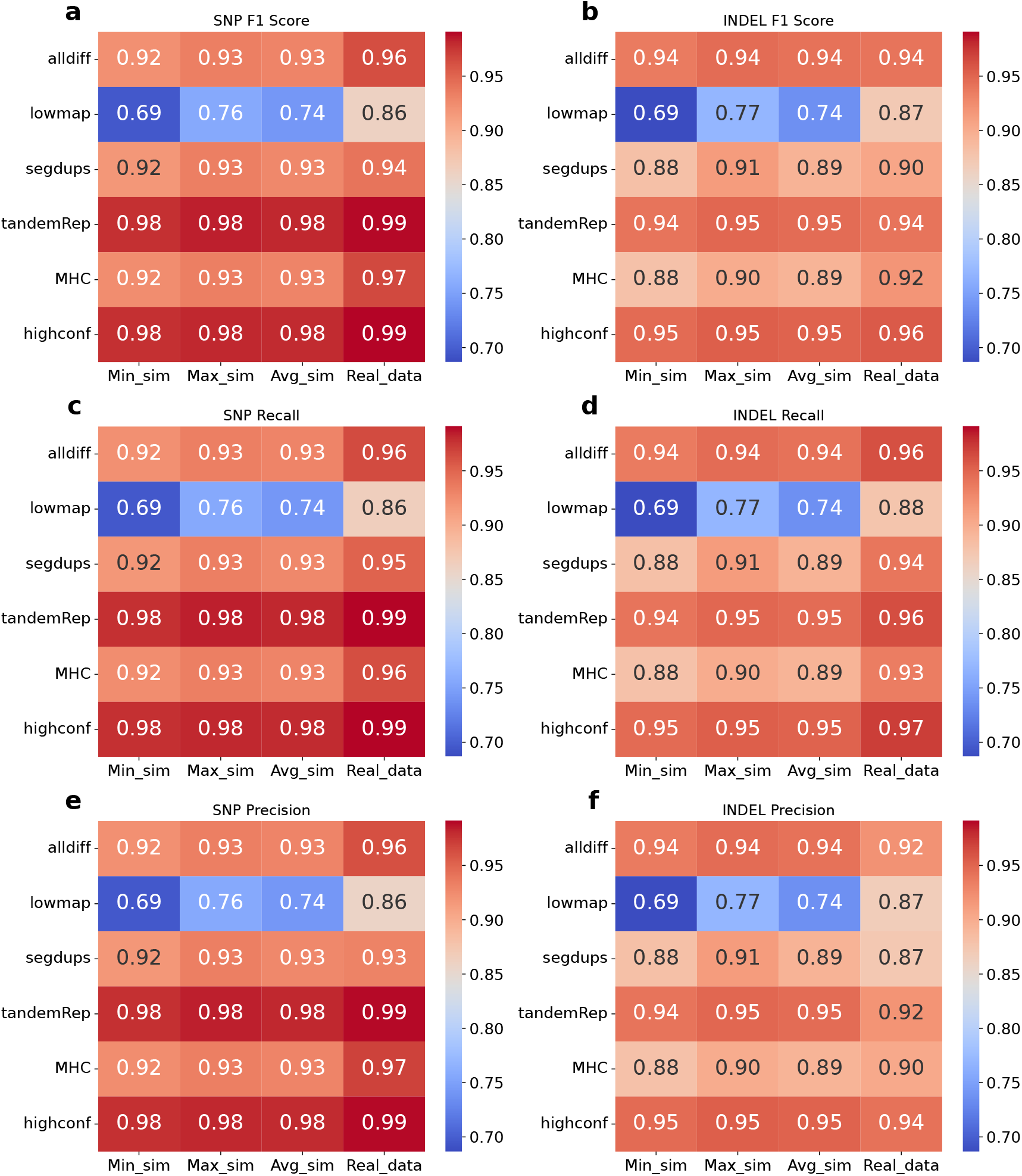
Simulated and real PE100_stLFR SNP INDEL calling performance comparison. *hap*.*py* v0.3.15 was applied to benchmark the SNP and small INDEL callset against the NIST v4.2.1 call set. Across 12 different simulation configurations, the minimum (Min sim), maximum(Max sim) and average(Avg sim) value of F1, recall and precision score were calculated. The evaluation was applied to different regions on the genome, including all difficult regions(alldiff), low mappability regions(lowmap), segmental duplication regions(segdups), tandem repeat regions(tandemRep), major histocompatibility complex regions (MHC) and high confidence region defined by GIAB for HG002(highconf). **a**. F1 for SNP calling. **b**. F1 for INDEL calling. **c**. Recall for SNP calling. **d**. Recall for small INDEL calling. **e**. Precision for SNP calling. **f**. Precision for small INDEL calling.

Overall, for SV calling, the simulated data exhibited similar trade-off patterns for both insertion and deletion SVs compared with real stLFR data. Further, our simulated data demonstrated SNP and small INDEL calling performance closely matching that of real stLFR data. These results suggested that our simulation framework realistically captured the characteristics of stLFR sequencing data. Therefore, we could confidently extend our analysis to other sequencing configurations, such as SE500_stLFR and SE1000_stLFR.

### Long single-end barcoded reads harbor strong potential for SV calling

In this section, we evaluated the performance of SV calling across three types of linked-reads libraries: PE100_stLFR, SE500_stLFR, and SE1000_stLFR. Each was sequenced at 35x coverage. For each library type, we analyzed 12 datasets to investigate the effects of sequencing parameters on SV calling. SVs were identified using Aquila stLFR (v2), an extension of the previous linked-read-based SV detection tool Aquila stLFR [9]. We benchmarked the resulting call sets against the GIAB SV benchmark set using Truvari (v4.0.0) [23] with the following parameters: *p*=0.5, *P* =0.5, *r*= 500, *O*= 0.01. Our evaluation focused on insertion SVs (INSs) and deletion SVs (DELs) larger than 50 bp within the high-confidence regions defined by GIAB.

For insertion SVs, the performance of the three library types varied significantly across the evaluated metrics, as shown in Figure 3a, c, and e. SE1000_stLFR demonstrated the highest F1 score, ranging from 0.81 to 0.86 with an average of 0.84 across all 12 datasets, indicating a superior balance between precision and recall. SE500_stLFR followed closely with F1 scores ranging from 0.78 to 0.83 (average 0.80), while PE100_stLFR exhibited the lowest performance, with F1 scores between 0.67 to 0.72 (average 0.70). For recall, SE1000_stLFR again outperformed the others, achieving values between 0.75 and 0.83 (average 0.82), reflecting its ability to detect true insertion SVs. SE500_stLFR showed moderate recall, ranging from 0.69 to 0.76 (average 0.73), whereas PE100_stLFR performed the worst, with recall values between 0.55 and 0.60 (average 0.58), highlighting its limitations in sensitivity. In contrast, precision was highest in SE500_stLFR, ranging from 0.87 to 0.92 with an average of 0.89, reflecting its strong accuracy in identifying true positives. PE100_stLFR followed closely, with a precision range of 0.84 to 0.91 (average 0.88), while SE1000_stLFR, despite its overall superior performance, showed a slightly narrower range of 0.87 to 0.89 (average 0.88). These results suggested that longer read lengths, as in SE1000_stLFR, enhanced recall but incurred a modest trade-off in precision, whereas shorter reads, such as PE100_stLFR, prioritized precision but struggled with recall.

**Figure 3.**
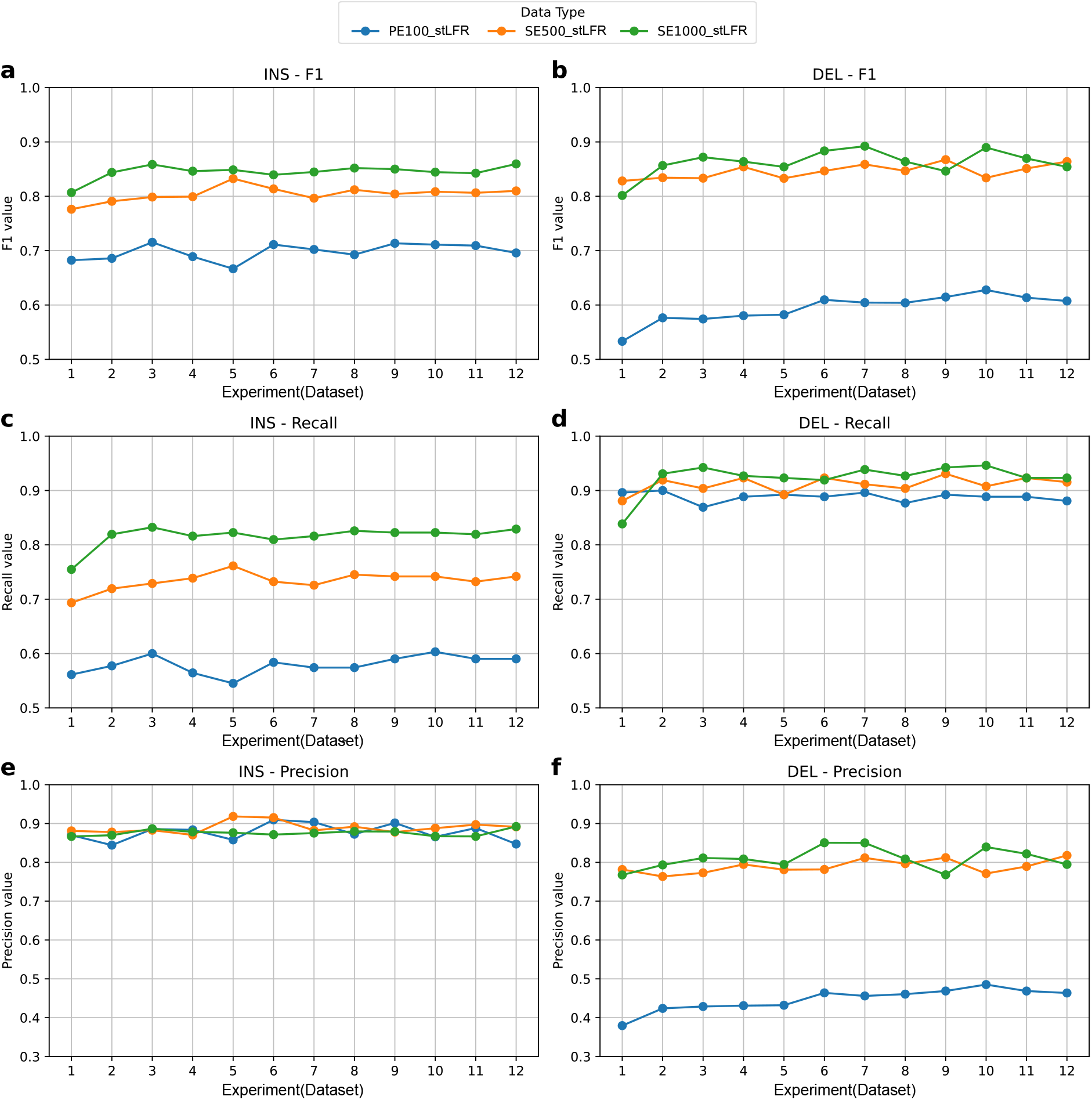
Structural variant detection performance across different single libraries. Aquila stLFR (v2) was applied to generate the SV callsets. Truvari (v4.0.0) was used to benchmark the result against GIAB SV benchmark set (parameters: *p*=0.5, *P* =0.5, *r*=500, *O*=0.01). Libraries are color-coded as follows: blue for PE100_stLFR, orange for SE500_stLFR, and green for SE1000_stLFR. **a**,**c**,**e** show the F1 score, recall, and precision for insertion SV (INS) calls **b**,**d**,**f** show the F1 score, recall, and precision for deletion SV (DEL) calls.

For deletion SVs, SE1000_stLFR consistently outperformed other libraries across most metrics (Figure 3b, d, and f). SE1000_stLFR achieved F1 scores ranging from 0.80 to 0.89, with an average of 0.86, indicating strong overall performance. SE500_stLFR was closely competitive, with F1 scores between 0.83 and 0.87 (average 0.85), while PE100_stLFR lagged behind, ranging from 0.53 to 0.63 (average 0.59), reflecting reduced balance between precision and recall. In terms of recall, all three datasets performed relatively well: SE1000_stLFR ranged from 0.84 to 0.95 (average 0.92), SE500_stLFR from 0.88 to 0.93 (average 0.91), and PE100_stLFR from 0.87 to 0.90 (average 0.89). These results collectively highlighted the robustness of DEL detection across datasets, with SE1000_stLFR showing the most consistent and accurate performance. Precision, however, remained a key differentiator. SE1000_stLFR again led, with precision values ranging from 0.77 to 0.85 (average 0.81), followed closely by SE500_stLFR, which ranged from 0.76 to 0.82 (average 0.79). In contrast, PE100_stLFR showed substantially lower precision, with values between 0.38 and 0.49 (average 0.45), highlighting a higher rate of false positives in deletion SV calls. These results emphasized that SE1000_stLFR, benefiting from longer reads, consistently offered the best trade-off between precision and recall. Conversely, the shorter reads of PE100_stLFR appeared to hinder accurate deletion SV detection, particularly in maintaining precision - likely due to the challenges of resolving complex genomic regions with limited sequence context.

Overall, insertion SV calls favored high precision but lower recall, while deletion SV calls showed higher recall at the cost of precision, emphasizing distinct tradeoffs between the two SV types. Longer reads were more effective in balancing these metrics, particularly for deletion SV detection. In contrast, shorter reads such as PE100_stLFR struggled, exhibiting reduced recall for insertion SVs and diminished precision for deletion SVs.

### Long single-end barcoded reads achieve comparable SV calling performance with long read based methods

Given the substantial improvement of SE1000_stLFR over PE100_stLFR in the preceding analyses, we next investigated how SE1000_stLFR compared with a broader range of sequencing-based strategies for structural variant detection, particularly long-read approaches that have shown strong performance in SV discovery. To address this, we selected SE1000_stLFR EXP7, a simulation setting that showed comparatively balanced performance across evaluation metrics, and benchmarked it on chromosome 6 of the HG002 SV Benchmark against representative methods spanning conventional short-read SV calling, pangenome-based short-read genotyping, and long-read sequencing. Specifically, we included Manta [25] as a representative short-read SV caller, PanGenie [26] as a pangenome-based short-read genotyping approach, and VolcanoSV [19] as a long-read–based SV calling benchmark. VolcanoSV was applied to PacBio HiFi data, whereas PanGenie genotyped Illumina short reads against a pangenome graph built from nine diploid high-quality assemblies. This comparison enabled us to place SE1000_stLFR in the context of both conventional short-read analysis and more advanced graph-assisted or long-read–enabled SV detection frameworks.

As summarized in Table 3, SE1000_stLFR showed consistently strong performance for both insertion and deletion SVs. Overall, it substantially outperformed the conventional short-read caller Manta, achieved performance comparable to PanGenie, and approached that of VolcanoSV, which remained the strongest method overall. For insertion SVs, SE1000_stLFR delivered a more favorable balance between precision and recall than Manta and surpassed PanGenie in overall accuracy, although VolcanoSV retained the best performance. For deletion SVs, SE1000_stLFR again clearly exceeded Manta and remained slightly stronger than PanGenie in overall detection performance, while trailing VolcanoSV only modestly. Genotyping concordance showed a similar overall trend, although with some differences across variant classes. SE1000_stLFR maintained strong insertion SV genotyping performance, whereas its deletion SV genotype concordance was lower than that of the other methods despite its strong deletion SV detection accuracy. This suggests that, for SE1000_stLFR, residual limitations are more evident in genotype assignment than in SV discovery itself. Together, these results indicate that SE1000_stLFR markedly advances beyond conventional short-read SV calling and reaches performance competitive with pangenome-based short-read genotyping, while narrowing the gap to long-read– based SV detection. These findings support extendedlength single-end barcoded sequencing as a practical intermediate strategy between standard short-read and long-read technologies for structural variant analysis.

**Table 3.**
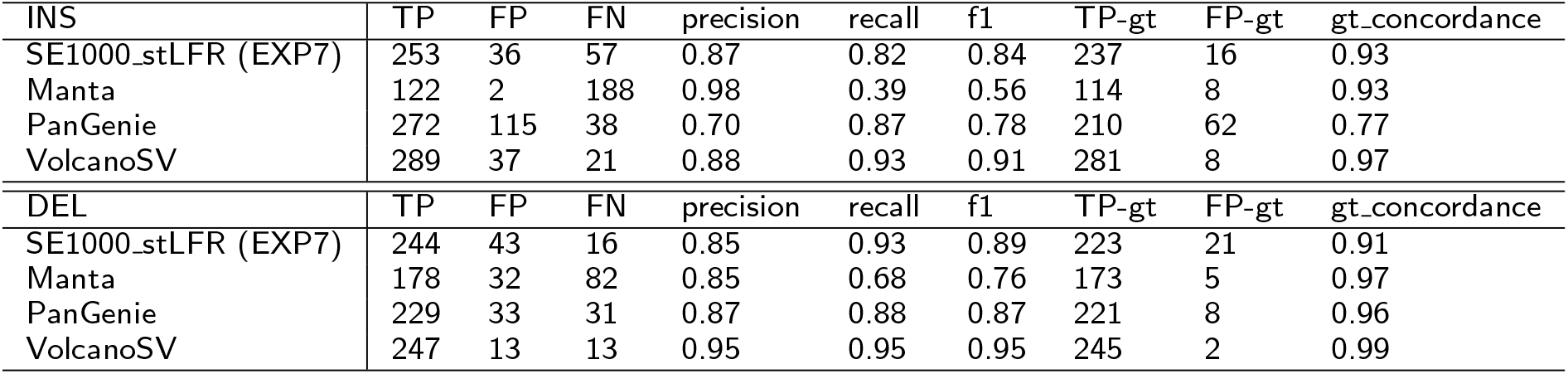
Comparison of structural variant calling performance for deletion and insertion SVs on chromosome 6 among SE1000_stLFR, Manta, PanGenie and VolcanoSV. Performance is summarized using true positives (TP), false positives (FP), false negatives (FN), precision, recall, F1 score, genotype true positives (TP-gt), genotype false positives (FP-gt), and genotype concordance. Across both deletion and insertion SV detection, SE1000_stLFR achieved competitive performance relative to the long-read-based caller VolcanoSV and the pangenome-based genotyper PanGenie, indicating that 1000-bp single-end barcoded reads can recover much of the SV calling capability typically associated with long-read approaches.

| INS | TP | FP | FN | precision | recall | f1 | TP-gt | FP-gt | gt_concordance |
| --- | --- | --- | --- | --- | --- | --- | --- | --- | --- |
| SE1000_stLFR (EXP7) | 253 | 36 | 57 | 0.87 | 0.82 | 0.84 | 237 | 16 | 0.93 |
| Manta | 122 | 2 | 188 | 0.98 | 0.39 | 0.56 | 114 | 8 | 0.93 |
| PanGenie | 272 | 115 | 38 | 0.70 | 0.87 | 0.78 | 210 | 62 | 0.77 |
| VolcanoSV | 289 | 37 | 21 | 0.88 | 0.93 | 0.91 | 281 | 8 | 0.97 |
| DEL | TP | FP | FN | precision | recall | f1 | TP-gt | FP-gt | gt_concordance |
| SE1000_stLFR (EXP7) | 244 | 43 | 16 | 0.85 | 0.93 | 0.89 | 223 | 21 | 0.91 |
| Manta | 178 | 32 | 82 | 0.85 | 0.68 | 0.76 | 173 | 5 | 0.97 |
| PanGenie | 229 | 33 | 31 | 0.87 | 0.88 | 0.87 | 221 | 8 | 0.96 |
| VolcanoSV | 247 | 13 | 13 | 0.95 | 0.95 | 0.95 | 245 | 2 | 0.99 |

## Discussion

In this study, we evaluated the capabilities and limitations of long single-end barcoded reads of 500 bp (SE500_stLFR) and 1000 bp (SE1000_stLFR), a conceptual extension of the traditional paired-end linked-read technology (PE100_stLFR), in detecting structural variants (SVs).

In terms of structural variant detection, our analysis underscores the significant impact of read length on SV calling, clearly demonstrating that longer single-end barcoded reads substantially enhance SV detection accuracy compared to shorter paired-end linked-reads. Specifically, SE1000_stLFR consistently provided the optimal balance between precision and recall, enabling robust identification of insertion and deletion SVs greater than 50 bp. SE500_stLFR demonstrated intermediate performance, falling between SE1000_stLFR and PE100_stLFR. While PE100_stLFR reads yielded high precision in insertion SV detection and high recall in deletion SV detection, their limited length restricted the recall in insertion SV detection and precision in deletion SV detection, resulting overall lower accuracy. Further, our analysis revealed that SV discovery using SE1000_stLFR reads achieved performance comparable to long-read–based approaches while remaining substantially more cost-effective. These findings highlight that increasing read length in linked-read sequencing markedly enhances the resolution of structural complexity, helping to overcome limitations inherent to conventional short-read sequencing methods. The strong performance and low cost of long singleend barcoded reads therefore make them a promising and cost-effective alternative to more expensive longread technologies. Employing longer single-end reads, or strategically integrating them in hybrid approaches, represents a promising direction for improving comprehensive genomic analyses and structural variant detection in future designs of linked-read sequencing libraries and studies.

## Conclusion

In this study, we systematically investigated the impact of different linked-read sequencing designs on structural variant (SV) detection. Our results show that long single-end barcoded reads, particularly SE1000_stLFR, substantially improve SV detection by achieving a more favorable balance between precision and recall. Overall, this work highlights the potential of long single-end barcoded reads as a powerful extension of existing stLFR technology: even a modest increase in read length can yield substantial gains in both detection performance and cost efficiency. If such long single-end barcoded reads become technically feasible, they could significantly advance SV discovery by combining high accuracy with a cost-efficient sequencing design.

## Declarations

### Ethics approval and consent to participate

This study is based exclusively on simulated sequencing data and publicly available reference genomes. It does not involve human participants, animals, or identifiable personal data. Therefore, ethical approval and consent to participate were not required.

### Consent for publication

Not applicable.

### Availability of data and materials

All VCF files supporting the findings of this study are available at https://doi.org/10.5281/zenodo.14338785.

The difficult genomic regions analyzed in this study were obtained from the NIST v4.2.1 genome stratifications v3.1 dataset, including all difficult regions (https://ftp-trace.ncbi.nlm.nih.gov/ReferenceSamples/giab/release/genome-stratifications/v3.1/GRCh37/Union/GRCh37_alldifficultregions.bed.gz), low-mappability regions (https://ftp-trace.ncbi.nlm.nih.gov/ReferenceSamples/giab/release/genome-stratifications/v3.1/GRCh37/mappability/GRCh37_lowmappabilityall.bed.gz), segmental duplication regions (https://ftp-trace.ncbi.nlm.nih.gov/ReferenceSamples/giab/release/genome-stratifications/v3.1/GRCh37/SegmentalDuplications/GRCh37_segdups.bed.gz), tandem repeat regions (https://ftp-trace.ncbi.nlm.nih.gov/ReferenceSamples/giab/release/genome-stratifications/v3.1/GRCh37/LowComplexity/GRCh37_AllTandemRepeatsandHomopolymers_slop5.bed.gz), and the MHC region (https://ftp-trace.ncbi.nlm.nih.gov/ReferenceSamples/giab/release/genome-stratifications/v3.1/GRCh37/OtherDifficult/GRCh37_MHC.bed.gz). NIST v4.2.1 HG002 GRCh37 call set is available at https://ftp-trace.ncbi.nlm.nih.gov/ReferenceSamples/giab/release/AshkenazimTrio/HG002_NA24385_son/NISTv4.2.1/GRCh37/

HG002 Illumina sequencing reads used to benchmark PanGenie and Manta were obtained from https://www.ebi.ac.uk/ena/browser/view/PRJEB35491. The assemblies used to build the pangenome graph for Pan-Genie are available at https://s3-us-west-2.amazonaws.com/human-pangenomics/working/HPRC_PLUS/SAMPLE/assemblies/year1_freeze_assembly_v2/, where SAMPLE should be replaced with the corresponding sample name. SV calling results generated by VolcanoSV were obtained from https://zenodo.org/records/10456758.

### Code availability

stLFR-sim is available at https://github.com/maiziezhoulab/stLFRsim under the MIT License.

### Author contributions

X.M.Z. and B.A.P. conceived and led this work. C.L., Y.H.L., Z.Z and L.Z. designed the framework. C.L., Y.H.L. and H.L. implemented the framework and performed all analysis. C.L., Y.H.L., and X.M.Z. wrote the manuscript. All authors reviewed the manuscript.

## Acknowledgements

Not applicable

